# An inter-chromosomal transcription hub activates the unfolded protein response in plasma cells

**DOI:** 10.1101/295915

**Authors:** Alexandra Bortnick, Zhaoren He, Megan Aubrey, Vivek Chandra, Matthew Denholtz, Kenian Chen, Yin C Lin, Cornelis Murre

**Author notes:** These authors made equal contributions (Alexandra Bortnick, Zhaoren He). Correspondence should be addressed to C.M.

## Abstract

Previous studies have indicated that the transcription signature of antibody-secreting cells is closely associated with the induction of the unfolded protein response pathway (UPR). Here we have used genome-wide and single cell analyses to examine the folding patterns of plasma cell genomes. We found that plasma cells adopt a cartwheel configuration and undergo large-scale changes in chromatin folding at genomic regions associated with a plasma cell specific transcription signature. During plasma cell differentiation, Blimp1 assembles into an inter-chromosomal transcription hub with genes associated with the UPR, biosynthesis of the endoplasmic reticulum (ER) as well as a cluster of genes linked with Alzheimer’s disease. We suggest that the assembly of the Blimp1-UPR-ER transcription hub permits the coordinate activation of a wide spectrum of genes that collectively establish plasma cell identity.

---

The genome is folded into chromosome territories that, with the exception of nucleoli, rarely intermingle^1,2^. Chromosomes themselves are organized as tandem arrays of loops that intermingle to establish euchromatic (A) or heterchromatic (B) compartments^3^. While the euchromatic compartment is predominantly positioned in the nuclear interior, a large fraction of the heterochromatic compartment is associated with the nuclear lamina^4,5^. During developmental progression, regulatory and coding elements often reposition from the lamina to the nuclear interior and vice versa to modulate patterns of gene expression^6,7^. An additional layer of chromatin architecture involves the assembly of nuclear bodies such as nucleoli and perinucleolar heterochromatin, which, similar to the nuclear lamina, have been associated with inactive gene transcription^8,9^.

Plasma cells are terminally differentiated B cells responsible for maintaining protective serum antibody titers that serve as the first line of defense against infection^10^. Plasma cells are also a source of pathogenic auto-antibodies, and in malignancy, the origin of plasmacytomas, light chain amyloidosis, monoclonal gammopathies, and multiple myeloma^11,12^. In response to pathogens, naïve B cells (also referred to as interphase, mature, resting, or quiescent) rapidly proliferate to form a population of activated B cells, which further differentiate into germinal center B cells or extra-follicular antibody-secreting plasma cells (also referred to as plasmablasts). Recent experiments tracking antigen-specific responses over hundreds of days have revealed plasma cell subsets with varying lifespans^13,14^. These findings raise questions about the minimum genomic requirements to establish plasma cell identity. Dramatic morphological and transcriptional changes accompany B cell activation and differentiation, many initiated within hours of stimulation ^13-15^. Studies linking these changes to chromatin reorganization events have focused on activated B cells (24 hours after stimulation), germinal center B cells, or mixed populations of activated and differentiated cells. ^16,17,18,19^

B cell development relies on a highly regulated and hierarchical program of gene expression. In early B cells, E-protein E2A induces the expression of FOXO1 and early B cell factor 1 (EBF1), which in turn activates the expression of FOXO1 in a feed-forward loop, ultimately activating PAX5 expression. Terminal B cell differentiation extinguishes this early signature while activating factors such as BLIMP1 and XBP1^20,21^. Repression of early B cell genes is central to initiation of the plasma cell program as forced expression of PAX5 interferes with antibody secretion^22^. In contrast, sustained expression of E-box transcription factors E2A (*Tcf3*) and E2-2 (*Tcf4*) are crucial to the plasma cell program^23-25^.

Plasma cells also constitutively activate the unfolded protein response (UPR), a specializing sensing mechanism for detecting and processing large amounts of protein shuttled through the endoplasmic reticulum (ER). The three sensors known to implement the UPR pathway are inositol-requiring enzyme 1 (IRE 1), PKR-like ER kinase (PERK), and activating transcription factor 6 (ATF6)^26^. When unfolded protein accumulates, ATF6 translocates the Golgi to drive ER quality control and capacity. IRE1 oligomerizes and activates ribonucleases, which splice x-box-binding protein 1 (XBP1) mRNA. XBP in turn upregulates chaperones and expands ER function and capacity. PERK oligomerizes and phosphorylates eukaryotic translation initiation factor 2 α (eIF2α). Phosphorylated eIF2α inhibits global protein translation while favoring translation of activating transcription factor 4 (ATF4) mRNA. ATF4 increases ER capacity as well as strongly induces C/EBP homologous protein (CHOP). If ER stress is not resolved through the UPR, CHOP induces apoptosis by inhibition of Bcl-2 and induction of Bim. XBP-1 null B cells are unable to secrete Ig or fully differentiate to plasma cells^27^. In differentiated plasma cells, the transcriptional regulator *Prdm1* (encodes for BLIMP-1) was shown to regulate the expression of *Xbp1*, *Atf4* and *Atf6* as well other downstream components of all three pathways^28^. In other work, plasma cell differentiation was found to proceed via a non-canonical PERK pathway^29,30^ where *Atf4* was important for optimal antibody secretion^31^ and plasma cell survival^32^ via an atypical activation of CHOP^33^.

Despite their essential role in health and disease, a description of the chromatin topology and linear genomic features of plasma cells remains to be determined^34^. Here, we mapped architectural changes during plasma cell differentiation. We found that the plasma cell genomic architecture is unique among cells in the B lineage. Global analysis in conjunction with modeling revealed that the genomes of antibody-secreting cells were enriched for chromosomes that adopted elongated and helical structures. Plasma cells also generated an intricate network of inter-chromosomal interactions involving *Prdm1* and genes essential for the unfolded protein response (UPR) and the assembly of the endoplasmic reticulum,. Together, these findings reveal how during plasma cell differentiation chromosomes adopt a unique chromosomal architecture to assemble an inter-chromosomal UPR hub to coordinately activate the expression of genes that specify plasma cell fate.

## RESULTS

### Plasma cell genomes are depleted for long-range intra-chromosomal interactions

As a first approach to determine how the three-dimensional organization of the plasma cell genome differs from that of naïve B cells, we performed genome-wide tethered chromosomal conformation capture analysis (TCC) using purified primary mouse follicular B cells and plasma cells^35^. Specifically, follicular naïve B cells (CD23^+^) were isolated from mice that carry a GFP reporter allele within the *Prdm1* locus,^36^ which were then activated *in vitro* to generate plasma cells. As expected, the average intra-chromosomal contact probability as a function of genomic distance revealed that the TCC reads for both naïve B and plasma cells decay as a function of genomic distance (**Fig. 1a**). Notably, however, compared to naïve B cells, plasma cells were depleted for intra-chromosomal interactions separated more than 10 Mb but were enriched for intra-chromosomal interactions that spanned less than 10 Mb (**Fig. 1a**). Circos plots derived from contact maps associated with naïve and plasma cells confirmed enrichment for intra-chromosomal genomic interactions that primarily involved euchromatic regions as denoted by PC1^+^ regions (**Fig. 1b**).

**Figure 1.**
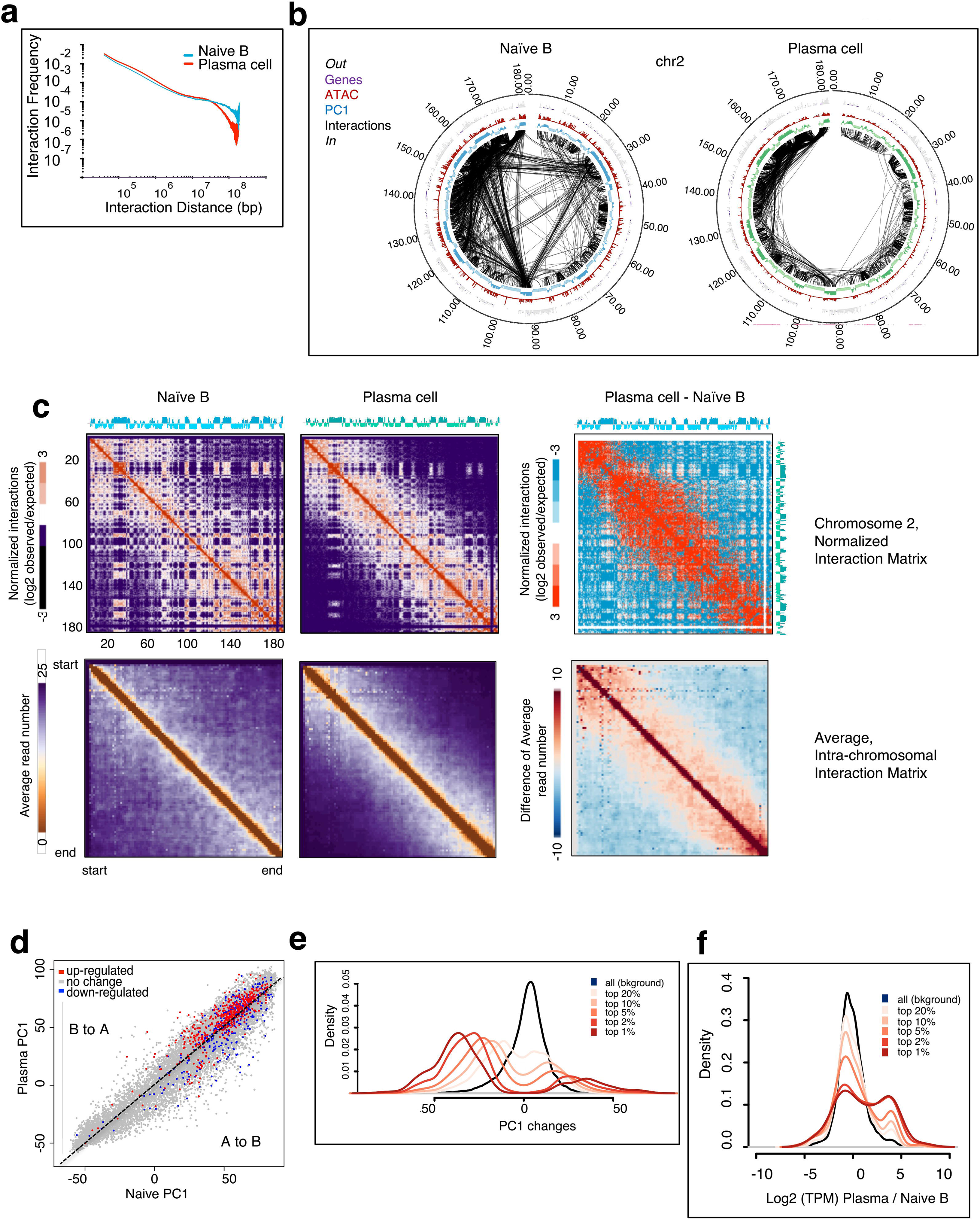
Plasma cell differentiation involves large-scale changes in intra-chromosomal interactions. **(a)** Intra-chromosomal interaction frequencies across the genome at 50 kb resolution for naïve B cells and plasma cells. **(b)** Circos plot of significant intra-chromosomal interactions within chromosome 2 for naïve B and plasma cells. Interactions with p-values less than 10^−9^ are shown. PC1, ATAC, and gene positions are shown. Bin size, 50 kb. Numbers at the margins indicate genomic position in megabases. **(c)** TCC interaction matrix for chromosome 2 (top) and for all the chromosomes (bottom) at 100 kb resolution for naïve B cells and plasma cells. The right panels indicate differential interaction matrices. Red represents enrichment of contact frequencies, whereas purple represents depletion of normalized interaction frequencies in plasma cells. **(d)** PC1 comparison between naïve B cells and plasma cells overlaying RNA-seq expression. Each spot represents a gene. PC1 values are calculated as the average PC1 of the 50 kb bins overlapping with the gene. RNA-seq change was calculated as the fold TPM change of the gene’s in plasma cells and naïve B cells. A 5-fold TPM increase is marked as upregulated (red), a 5-fold decrease is marked as downregulated (blue); otherwise, unchanged (gray). **(e)** Change in PC1 values as it relates to the correlation difference (corrDiff). Each line represents the distribution of PC1 differences between plasma cell and naïve B of the regions satisfying the criteria indicated. For example, “top 1%” denotes the regions with the top 1% lowest (most dissimilar) corrDiff values. All background means all regions in the genome were considered. **(f)** Change in gene expression between plasma cells and naïve B cells, shown as the ratio of the log2 of TPM, as it relates to corrDiff.

Next, we constructed heat maps for naïve and plasma cell genomes (**Fig. 1c**). Heat maps representing interaction frequencies for chromosome 2 revealed a striking loss of long-range genomic interactions in plasma cells when compared to naïve B cells (**Fig. 1c, upper panel**). The differences were particularly pronounced in differential heat maps constructed by subtracting contact frequencies in plasma cells versus naïve B cells (**Fig. 1c, right upper panel**). The contact matrices derived from naïve and plasma cells showed a similar distribution for other chromosomes as revealed by plotting the intra-chromosomal frequency average for all chromosomes (**Fig. 1c; lower panel**). Taken together, these observations indicate that differentiating plasma cells adopt a unique nuclear architecture associated with a depletion for long-range intra-chromosomal interactions.

### Large-scale changes in compartmentalization in differentiating plasma cells

Previous studies revealed that a wide spectrum of genomic regions switched compartments during the developmental transition from the pre-pro-B to the pro-B cell stage. To examine whether likewise coding and/or regulatory DNA elements reposition during plasma cell differentiation, we compared the PC1 values associated with genomic regions derived from TCC data for naïve B cells versus plasma cells **(Fig. 1d)**. During the naïve to plasma cells transition, 347 domains repositioned from the A to B compartment, whereas 166 domains repositioned from the B to A compartment. To directly compare compartments in naïve B cells and plasma cells, correlation difference (corrDiff) values were calculated. corrDiff directly measures the interaction profiles of a given locus between two experiments. As corrDiff approaches 1, two regions become more similar. To determine the compartment (A or B) associated with each corrDiff value, we compared the PC1 values of corrDiff low (dissimilar) regions. This analysis revealed that during the naïve to plasma cell transition, regions associated with the *Ebf1, Tcrb, Bcl6* and *Bc1lla* loci changed interaction profiles and switched from the transcriptionally permissive to the transcriptionally repressive compartment (**Supplementary Table 1)**. Additionally, an ensemble of genomic regions, including regions associated with the immunoglobulin heavy and light chains, *Ell2, Sec61g, CD36, Cdkn2a (p16INK4)* and *Cdkn2b (p15INK4b)*, which induce cell cycle arrest in G1, also changed interaction profiles and repositioned from the transcriptionally repressive to the permissive compartment during plasma differentiation (**Supplementary Table 1**).

**TABLE.** DNA FISH probes

| Region of interest | Chromosome | Start (bp) | End (bp) | BAC-PAC Identifier |
| --- | --- | --- | --- | --- |
| EBF1 promoter | Chromosome | 44416074 | 44457525 | WIBR1-0109P11 |
| EBF1 distal enhancer | Chromosome | 45305517 | 45347678 | WIBR1-0350 |
| Ebf1 super-enhancer | Chromosome | 44941681 | 45143731 | RP23-118P17 |

To examine whether changes in compartmentalization correlated with changes in gene expression, we plotted the ratio of RNA abundance in naïve B versus plasma cells to PC1 values (**Fig. 1d**). As expected, during the naïve to plasma cell transition, a significant fraction of genomic regions that switched from compartment A to B and vice versa showed coordinate changes in RNA abundance (**Fig. 1d**). Next, we clustered regions into quantiles based on their corrDiff value: top 1% most different correlation profiles between naïve B cells and plasma cells, top 2% most different correlation profiles, etc. (**Fig. 1e**). Of the regions with the top 1% most different correlation profiles during the plasma cell transition, many switched from euchromatin to heterochromatin as revealed by a greater read density of negative rather than positive PC1 values (**Fig. 1e**). Interestingly, most of the regions that switched to the heterochromatic compartment were already silenced in naive cells, as only modest declines in transcriptional silencing for these regions were observed **(Fig. 1f)**. In contrast, the regions that switched from heterochromatin to euchromatin were accompanied by substantial increases in transcription levels (**Fig. 1f**). Taken together, these data indicate that the acquisition of a transcription signature associated with antibody-secreting cells is closely associated with large-scale changes in compartmentalization. In plasma cells, genes silenced in naïve B cells repositioned to the heterochromatic compartment where they were further silenced whereas inactive genes relocating to the euchromatic compartment became highly transcribed.

### Large-scale changes in the chromatin folding and compartmentalization of genes that orchestrate plasma cell fate

The data described above indicate that plasma cell differentiation is closely associated with large-scale changes in chromatin folding and compartmentalization. The extent to which PC1 and corrDiff analyses predicted local interaction changes varied across loci. Thus, to confirm and extend the PC1 and corrDiff analyses, we examined whether genes known to facilitate plasma cell differentiation were linked with alterations in chromatin topology and compartmentalization. TCC contact maps and PC1 and corrDiff tracks across the *Irf4, Xbp1, Bcl6 and Pax5* loci were compared for naïve B versus Blimp-expressing plasma cells (**Figs. 2a-d, Supplementary Fig. 1**). To identify the enhancer repertoire associated with naïve B and plasma cells and long-range genomic interactions, sorted bone marrow plasma cells from Blimp-GFP mice were analyzed using ATAC-Seq (assay for transposase-accessible chromatin). The interaction pattern between the open regions and the Irf4 promoter changed during the plasma cell transition, largely due to the significant increase in the number of open regions across the locus in plasma cells (**Fig. 2a**). In contrast, the number ATAC-seq peaks around the *Xbp1* locus remained similar between cell types, yet plasma cells established new promoter-enhancer interactions between pre-existing open regions (**Fig. 2b**). We observed the inverse pattern for genes associated with declining mRNA levels in differentiating plasma cells. *Bcl6* and *Pax5* exhibited loss of open regions accompanied by weakened promoter-enhancer interactions in plasma cells (**Fig. 2c,d**). We further investigated the relationship between ATAC-seq peaks and interaction patterns for other loci, including *Atf4*, *Myc*, and *Foxo1* loci (**Supplementary Fig. 3**).

**Figure 2.**
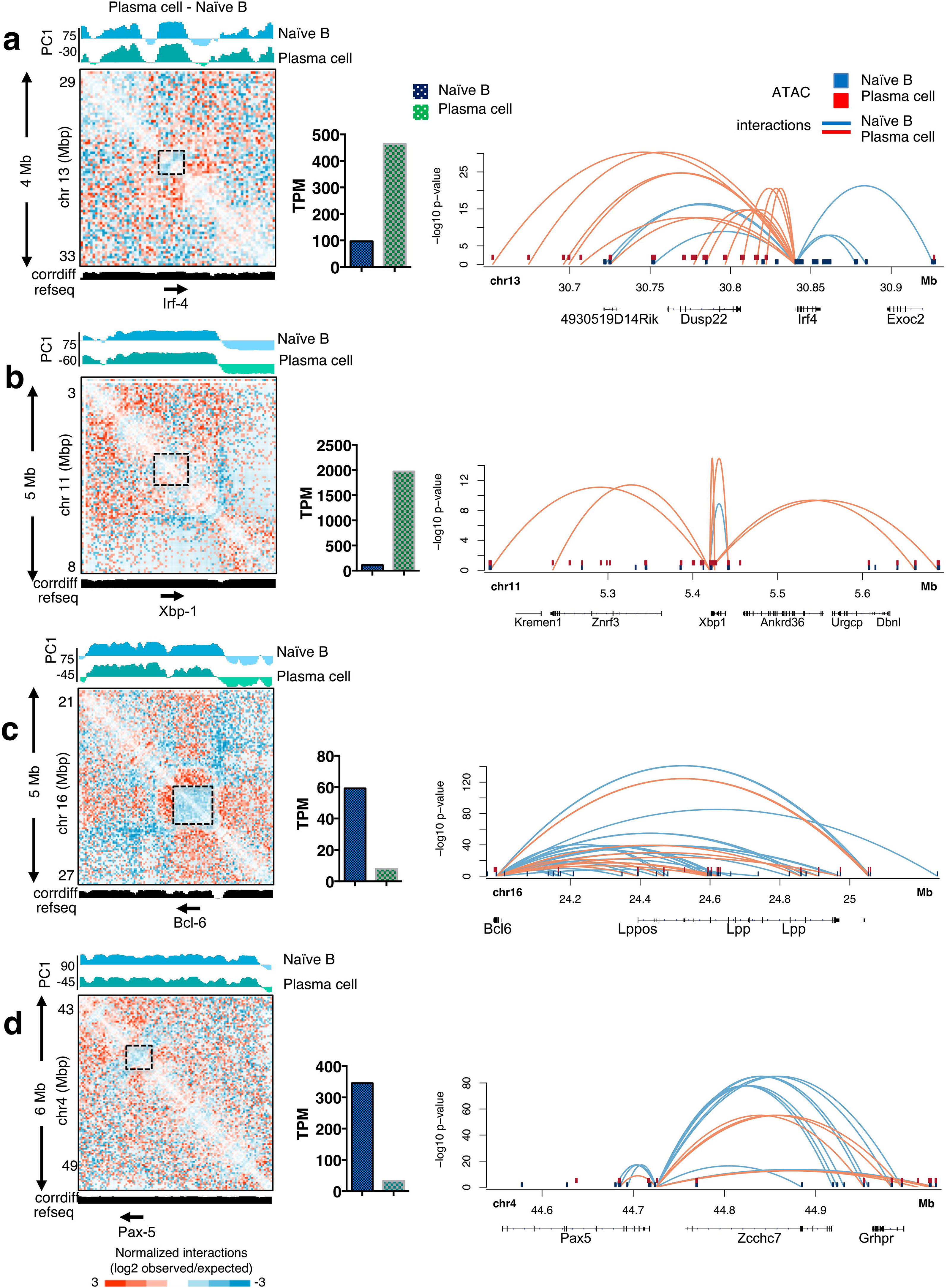
Plasma cell signature genes display changes in local interactions across enhancers and promoters coordinate with changes in gene transcription. **(a)** Contact map for chromosome 13 from 29 to 33 Mb at 100kb resolution (left). RNA level of *Irf4* in naïve B cells and plasma cells (middle). Sushi plot zooming in on the *Irf4* locus and associated ATAC peaks from 30.6 to 30.9 Mb (right). **(b)** Contact map for chromosome 11 from 3 to 8 Mb at 100kb resolution (left). RNA levels of *Irf4* in naïve B cells and plasma cells (middle). Sushi plot zooming in on the *Xbp1* locus from 51.5 to 56.9 Mb (right). **(c)** Contact map for chromosome 16 from 21 to 27 Mb at 100 kb resolution (left). RNA level of *Irf4* in naïve B cells and plasma cells (middle). Sushi plot zooming in on the *Bcl6* locus from 23.9 to 25.2 Mb (right). **(d)** Contact map for chromosome 4 from 43 to 49 Mb at 100 kb resolution (left). RNA level of *Irf4* in naïve B cells and plasma cells (middle). Sushi plot zooming in on the *Pax5* locus from 45.0 to 30.9 Mb (right).

**Figure 3.**
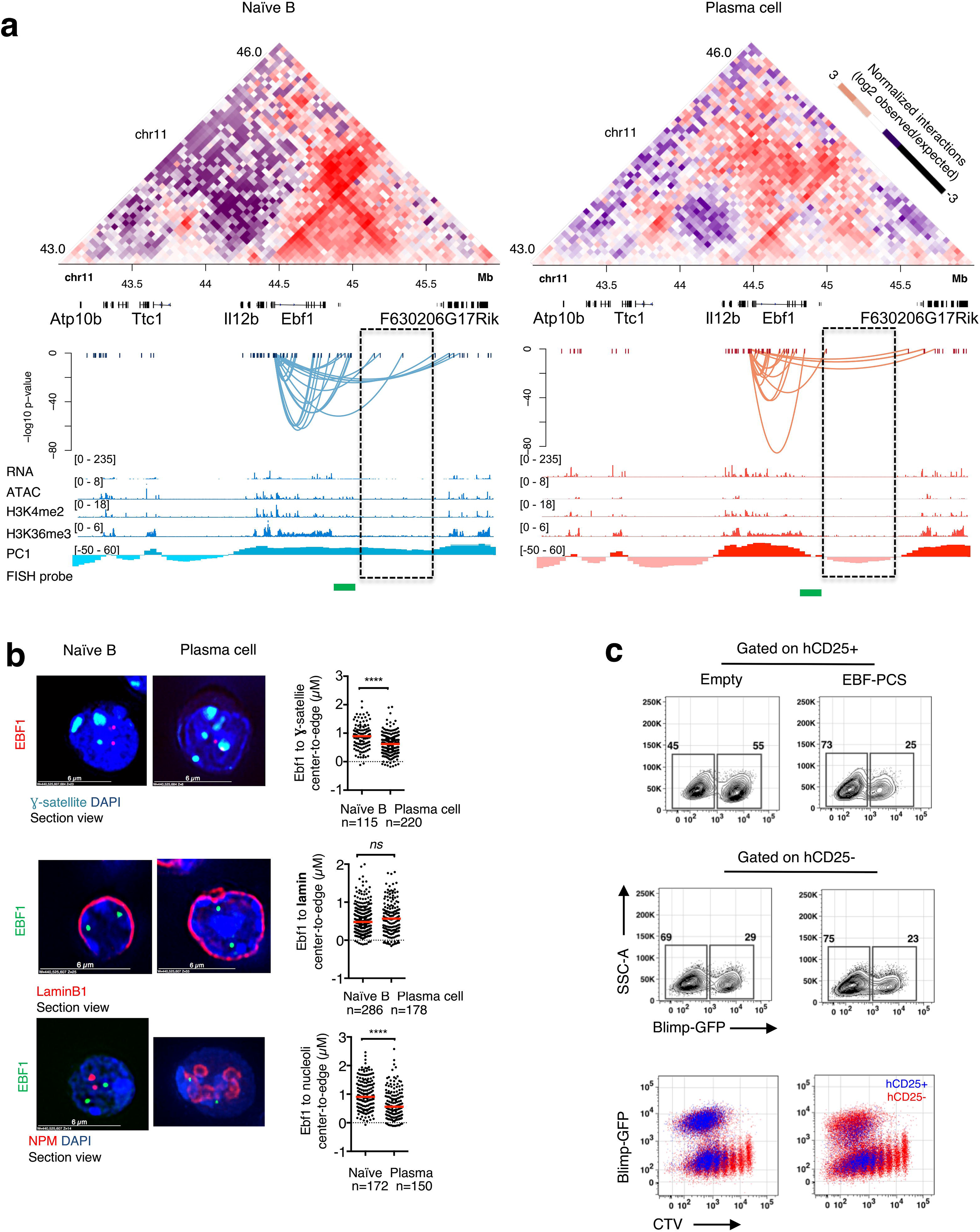
Ebf1 displays changes in local interactions and repositions away from the nuclear center towards repressive nuclear structures. **(a)** Contact map for chromosome 11 from 43 to 46 Mb at 100 kb resolution. Gene tracks from refseq. Sushi plot showing the interaction between Ebf1 promoter and the ATAC peaks in target region. IGV tracks of RNA-seq, ATAC-seq, H3K4me2, H3K36me3, and location of Ebf1 DNA-FISH probes (green) for naïve B cells (left) or plasma cells (right). **(b)** Example images from immunofluorescence for NPM (red) combined with DNA-FISH for the EBF1 locus (green). NPM foci were used to demarcate nucleoli (red) and DAPI was used to demarcate nuclear DNA (blue). **(c)** Representative flow plots of uninfected, empty vector, and EBF1-overexpressing activated B cells 48 hours after infection.

Previous studies demonstrated that the *Ebf1* locus is sequestered at the nuclear lamina in pre-pro-B cells but repositions to the nuclear interior in pro-B cells^6^. In contrast to the pre-pro-B to pro-B cell transition, where the *Ebf1* gene body switched compartments, we found that differentiating plasma cells instead only repositioned a distally located intergenic region from the A to the B compartment (**Fig. 3a**). To determine whether changes in the *Ebf1* locus were associated with changes in chromatin topology, genomic interactions across the *Ebf1* locus were compared for naïve B cells and Blimp-expressing plasma cells (**Fig. 3a)**. Compared to naïve B cells, differentiating plasma cells displayed declining levels of genomic interactions across the *Ebf1* locus, specifically across a cluster of distally-located enhancers marked by loss of ATAC-seq and H3K4me2 peaks (**Fig. 3a**). To validate that the *Ebf1* genomic region that switched from compartment A to compartment B during plasma cell differentiation repositioned, we used 3D-FISH in conjunction with antibodies that mark three repressive compartments, the nuclear lamina, nucleoli, and perinucleolar heterochromatin (“chromocenters”)^17,18^ (**Fig. 3b**). We found that in plasma cells the *Ebf1* locus rather than being sequestered at the nuclear lamina relocated towards heterochromatic regions associated with nucleoli and chromocenters, consistent with observations that the nucleolus, specifically its periphery, can act as an anchor for inactive chromatin^37^ (**Fig. 3b**). To confirm a potential inhibitory role for *Ebf1* in plasma cell differentiation, we overexpressed EBF1 in activated B cells and measured plasma cell differentiation by Blimp-1 expression 48 hours after infection. We found that EBF1 overexpression interfered with plasma cell differentiation, decreasing Blimp1^+^ cells by half in three day cultures (**Fig. 3c**). Taken together, these data indicate that the transition from the naïve B to the plasma cell stage is closely associated with the repositioning of genomic regions linked with the expression of developmental regulators that specify plasma cell identity.

### Plasma cell chromosomes adopt a cartwheel configuration

The data described above revealed that plasma cell differentiation was closely associated with changes in chromatin topology. As a first approach to determine how alterations in chromatin folding affect nuclear architecture, we built 3D-models for naïve and plasma cell chromosomes (**Methods**). Structures represent a statistical average of an ensemble of conformations from naïve B cells and plasma cells. Whereas naïve B cell chromosomes displayed spherical-like structures, plasma cell chromosomes adopted elongated and α-helical like structures (**Fig. 4a,b**). Remarkably, and consistent with previous electron microscopy^38^, the modeling of TCC reads derived from plasma cell genomes revealed a clock-face or cartwheel nuclear architecture (**Fig. 4c**) shaped by an “outer shell” primarily formed by the larger chromosomes (chrs. 1-9) (**Fig. 4d**). A rudimentary shell comprised of these chromosomes also appeared in naive B cells, but distinctly lacked the elongated chromosomes that established the cartwheel structure. This observation is consistent with previous studies in interphase cells, which also indicated that larger chromosomes tend to reside near the nuclear lamina^39^. In plasma cells, the elongated chromosomes showed polarity. Chromosomes aligned head-to-tail, situating the starts and ends of all chromosomes into closer proximity to one another (**Fig. 4e**). Consistent with previous imaging studies^19^, the most highly expressed gene loci in plasma cells, the immunoglobulin heavy and light chain loci, were predominantly positioned at the nuclear periphery (**Fig. 4f**). In naïve B cells, genomic regions associated with compartment A (transcriptionally permissive compartment) were enriched at the nuclear interior whereas many, but not all, of the PC1^+^ regions in plasma cells re-positioned outward towards the nuclear envelope (**Fig. 4g**). Inter-chromosomal interactions were mostly localized towards the nuclear interior of both naïve B cells and plasma cells (**Fig. 4h**). Finally, inter-chromosomal interactions were formed predominantly among PC1^+^ regions in both naïve B cells and plasma cells, although the distribution of inter-chromosomal interactions shifted slightly towards PC1^−^ regions in plasma cells (**Fig. 4i**). Taken together, these observations indicate that differentiating plasma cells are closely associated with a unique nuclear architecture enriched for elongated, helical chromosomes, which collectively generate a cartwheel-like structure.

**Figure 4.**
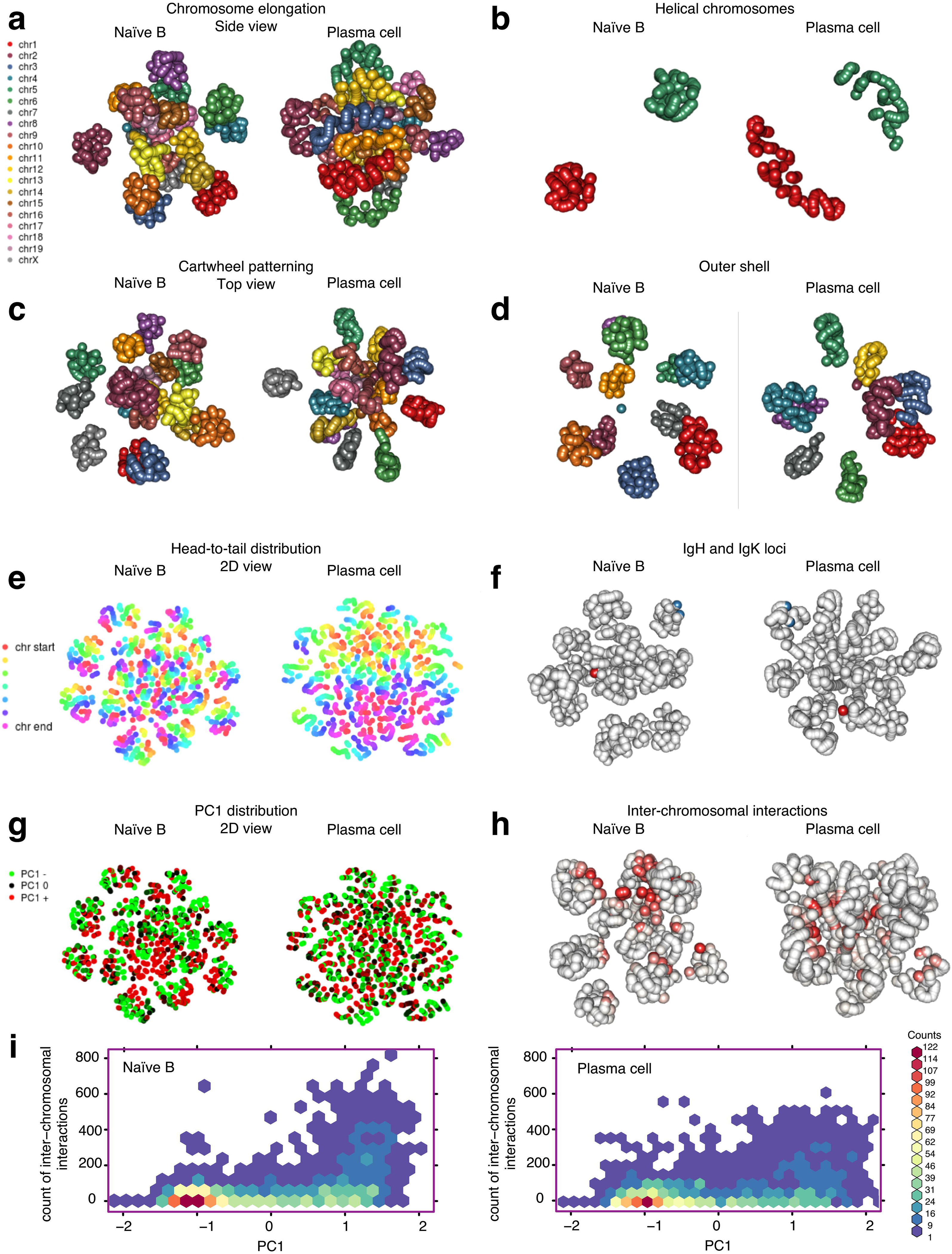
TCC-based modeling reveals defining features of the spatial arrangement and active regions of plasma cell chromosomes. Each dot represents a 2 Mb region. **(a)** Side view of all chromosomes, labeled in different colors to show the elongation of chromosomes in plasma cell. **(b)** Chromosomes 1 and 5 are shown as examples of the helical shape of chromosomes in plasma cells. **(c)** Top view of all chromosomes, showing the cartwheel pattern in plasma cells. **(d)** Outer shell of chromosomes. Chromosomes 1 to 9 in naïve B and chromosomes 1 to 8, 12, and 18 in plasma cells are shown. **(e)** Chromosome polarity in 2D view. All chromosomes are shown, colored as shown in legend to show the alignment of chromosome starts and ends in plasma cells. **(f)** Positioning of the IgH (chr. 12) and IgK (chr. 6) loci. All chromosomes displayed. **(g)** Positioning of the PC1^+^ (red) and PC1^−^ (green) regions for all chromosomes. **(h)** Inter-chromosomal interactions among all chromosomes. Greatest number of interactions (red) versus weakest number of interactions (pink) are displayed. **(i)** Heat map displaying the frequency of inter-chromosomal interactions for PC1^+^ and PC1^−^ regions. Counts are the number of regions in the genome at 1 Mb resolution. Interactive 3D plots are available online: http://bcell3d.ucsd.edu:3838/3Dsimulation_Bcell/

### Plasma cells adopt a spectrum of *de novo* inter-chromosomal interactions

The data described above indicate large-scale re-organization of intra-chromosomal genomic interactions in differentiating plasma cells. To examine whether and how plasma cell genomes differ from naïve B cells as it relates to intra-versus inter-chromosomal interactions, we plotted the total intra-chromosomal and inter-chromosomal paired-end tags for naïve B and plasma cells as a function of normalized chromosomal position (**Fig. 5a**). Notably, for large but not small chromosomes, plasma cells were severely depleted of long-range intra-chromosomal interactions at the chromosome end as compared to naïve B cells (**Fig. 1c and Fig. 5a, left panel**). In contrast, large chromosomes in plasma cells were associated with a significant increase in inter-chromosomal interactions at chromosome starts and ends (**Fig. 5a, right panel**). We next constructed inter-chromosomal heat maps for naïve and plasma cells (**Fig. 5b**). Consistent with the analysis described above, plasma cell genomes were substantially enriched for inter-chromosomal contact frequencies when compared to naïve B cells (**Fig. 5b, left**). Specifically, plasma cells were highly enriched for interactions involving the chromosome-ends between all pairs of chromosomes (**Fig. 5b, middle and right panels**). Interestingly, many inter-chromosomal interactions detected in plasma cells were pre-established in naïve B cells. Naïve B cells exhibited a scaffold of interactions that was preserved in plasma cells **(Fig. 5c; blue lines)**. In addition to these shared interactions, both naïve B cells and plasma cells displayed unique inter-chromosomal interactions. As observed in the contact matrices, the unique plasma cell interactions depicted in the Circos plots typically clustered toward chromosome ends whereas naïve B cell interactions spanned the chromosomes (**Fig. 5c; red lines**). Although both cell types contained similar numbers of significant inter-chromosomal interactions (p-value <10^−6^; 105,218 interactions in plasma cells; 101,564 interactions in naïve B cells), plasma cells displayed fewer strong interactions compared to naïve B cells (p-value <10^−20^; 9,244 interactions in plasma cells; 18,944 interactions in naïve B). The elongation and end-aligning of chromosomes in plasma cells appeared to not only weaken the pre-existing inter-chromosomal interactions (**Fig. 5d; upper panel)** but also facilitate only relatively weak new interactions (**Fig. 5d; lower panel**). Taken together, these observations indicate that differentiating plasma cells orchestrate an elaborate pattern of looping involving inter-chromosomal interactions often involving the large chromosomes.

**Figure 5.**
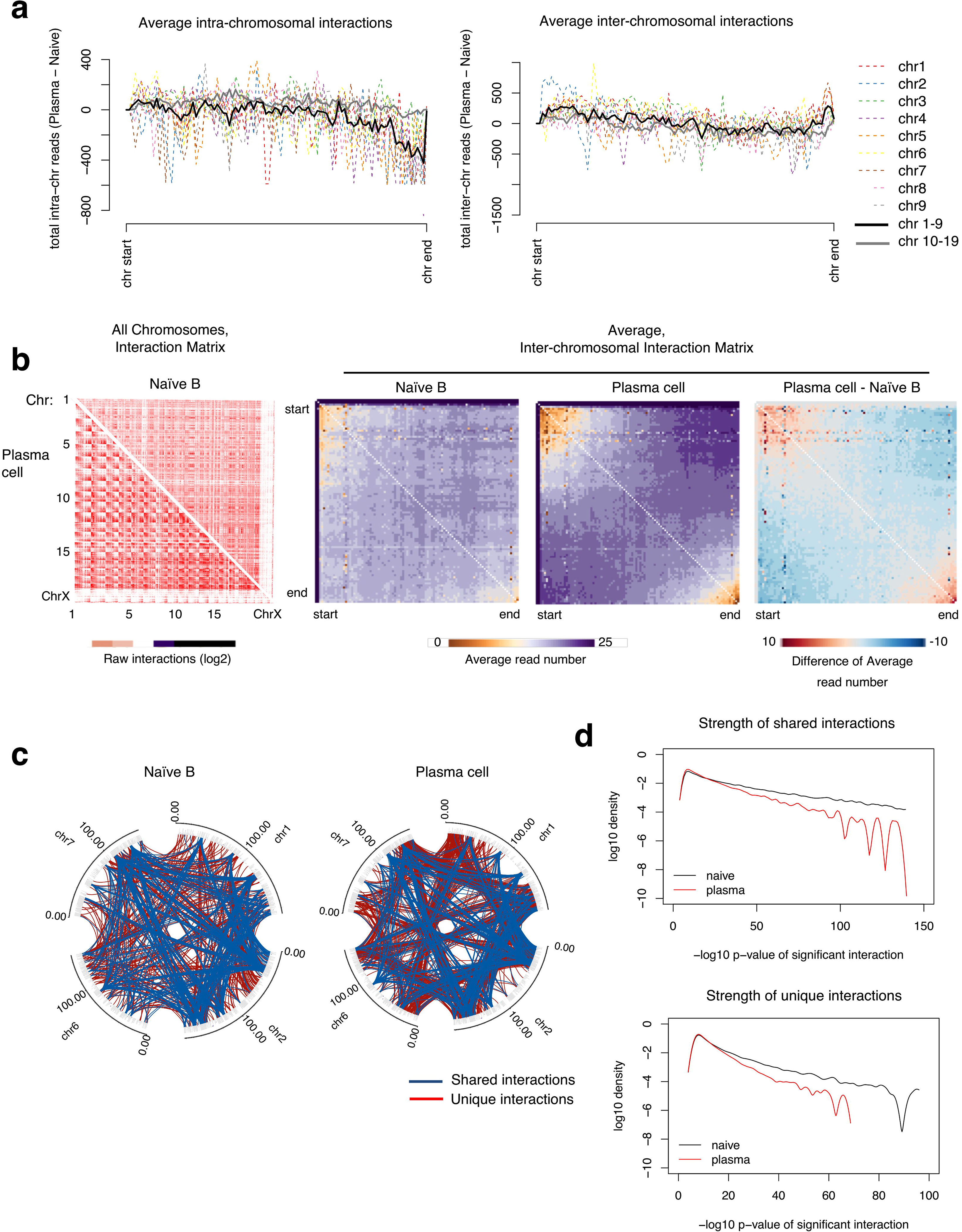
Plasma cell differentiation involves large-scale changes in inter-chromosomal interactions. **(a)** Intra- or inter-chromosomal TCC read number differences between plasma cells and naïve B cells. Average interactions of larger chromosomes 1-9 (black); average of smaller chromosomes 10-19 (gray). Chromosomes 1 to 9 are displayed as dotted lines as indicated by the legend. **(b)** Raw interaction matrix for naïve B and plasma cells (left); average inter-chromosomal interaction matrices (middle). Difference of the average inter-chromosomal interaction matrices between plasma and naïve cell (right). Red indicates gain in interaction; purple indicates depletion of interaction. **(c)** Circos plots showing significant inter-chromosomal interactions (p < 10^−4^) between chromosomes 1, 2, 6 and 7. Blue lines denote shared interactions between naïve B and plasma cells. Red lines indicate interactions unique to each cell type. **(d)** Distribution of log10 p-value of all significant inter-chromosomal interactions for naïve B cells and plasma cells separated by shared (top) versus unique (bottom) interactions.

### A transcriptional hub involving genes in the unfolded protein response pathway is assembled during plasma cell differentiation

To characterize in greater detail the ensemble of inter-chromosomal interactions associated with plasma cell differentiation and how they relate to compartmentalization and gene expression, we generated inter-chromosomal interaction cloud plots for naïve B and plasma cells (**Fig. 6a-d, Methods**). Inter-chromosomal cloud plots represent the interaction pattern of 1 Mb regions of the genome as a function of distance in 2D. For both naïve B cells and plasma cells, genomic regions enriched for inter-chromosomal interaction frequencies readily segregated into either compartment A or B (**Fig. 6a, top**). As expected, compartmentalization correlated well with gene expression levels for both naïve B and plasma cells (**Fig. 6a, middle**). Notably, we found that *Xbp1* and *Atf4*, two genes encoding for proteins involved in the UPR pathway, clustered in the inter-chromosomal interaction cloud within the euchromatic region (PC1>0) in both naïve B and plasma cells (**Fig. 6a, bottom**). In contrast, the relative spatial position of the *Prdm1* locus differed between naïve B cells and plasma cells, despite mild local intra-chromosomal interactions changes across the *Prdm1* locus (**Supplementary Fig. 3**). In naïve B cells, *Prdm1* had fewer significant interactions with the *Xbp1* and *Atf4* loci (**Fig. 6a**) whereas, in plasma cells, the *Prdm1* genomic region made more interactions with the *Xbp1* and *Atf4* loci (**Fig. 6a**). To validate this analysis, Circos plots were generated from the contact maps associated with naïve and plasma cells and gated on *Prdm1* (**Fig. 6b**). In addition to the *Xbp1* and *Atf4* interactions, the Circos diagrams revealed a spectrum of significant inter-chromosomal interactions unique to plasma cells (**Fig. 6b**). Using gene set enrichment analysis (GSEA) ^40^, we identified plasma cell-specific UPR-related genes interacting with the *Prdm1* locus, which not only included *Xbp1* and *Atf4* but also *Spcs1, Bax, Tmbim6,* and *Tmem129* (**Fig. 6c**). Additionally, we found significant enrichment for genes interacting with *Prdm1* in plasma cells involved in the physiology of the ER (57 genes), response to cellular stress (52 genes), and most prominently, Alzheimer’s disease (67 genes) (**Fig. 6c)**. The Alzheimer’s cluster was comprised of genes associated with metabolic processes, intracellular signal transduction, cell proliferation, and housekeeping genes. Inter-chromosomal interaction cloud analysis further revealed that the Alzheimer’s cluster of genes linked by inter-chromosomal interactions was pre-established in naïve B cells, which upon differentiation into plasma cells, connected with the *Prdm1* genomic region (**Fig. 6d**). Consistent with these findings, the enhancer repertoires of genes linked with Alzheimer’s disease as well as ER function were significantly enriched for BLIMP1 binding sites (**Fig. 6e**).

**Figure 6.**
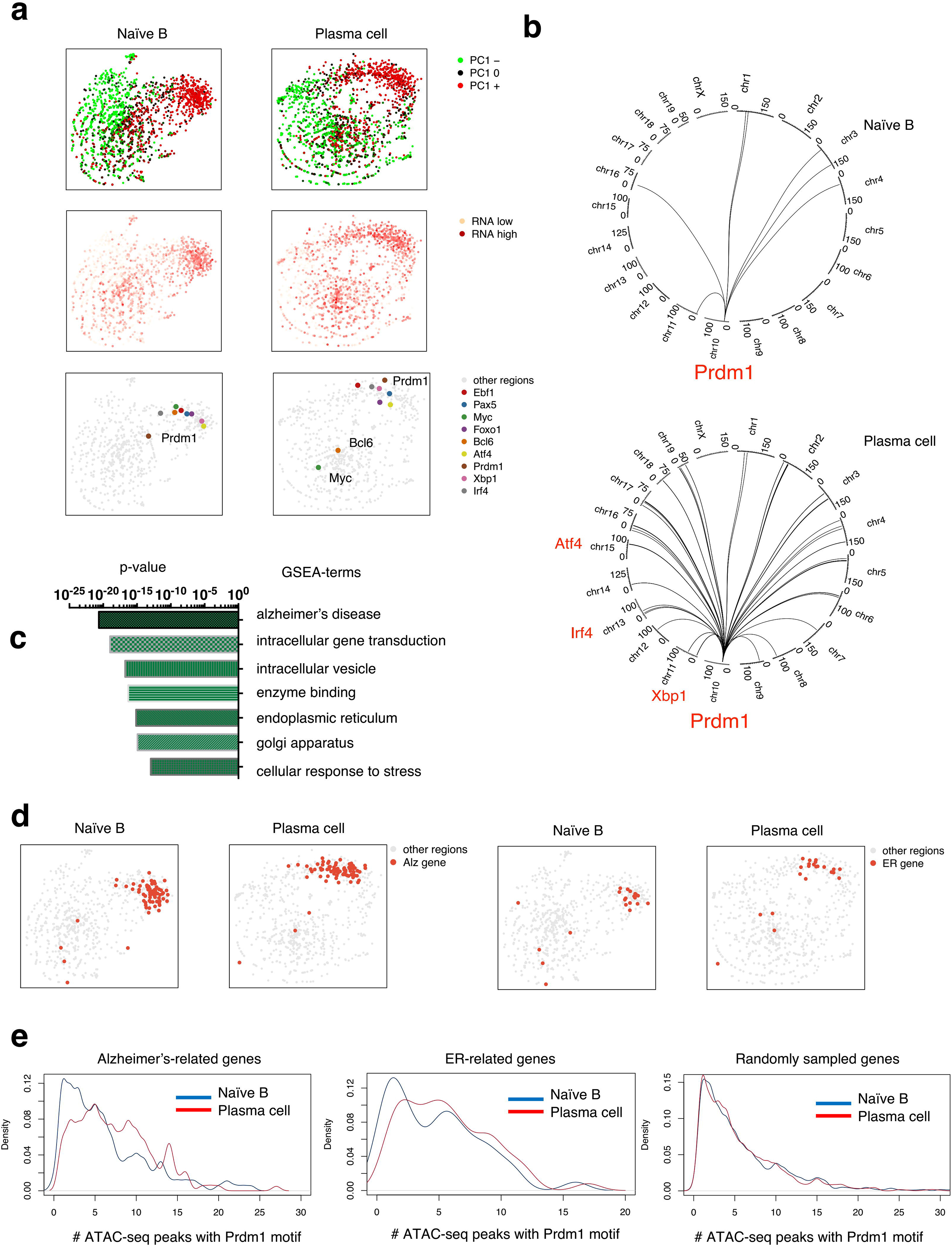
*Prdm1* forms inter-chromosomal interactions with ER and Alzheimer’s related genes. **a)** Inter-chromosomal interaction cloud plots showing the relative inter-chromosomal interaction preference of all genome regions (computed based on inter-chromosomal TCC reads). Each dot represents a 1 Mb region of the genome. The closer two dots in 2D distance, the more likely they share significant inter-chromosomal interactions. Dots are colored by PC1 value (top), RNA expression (middle) and genes of interest (bottom). **(b)** Circos plots for inter-chromosomal interactions centered on *Prdm1*. Genes contained within the 1 Mb interaction region and that appear in the inter-chromosomal cloud plot are indicated in red. **(c)** Gene set enrichment analysis (GSEA) performed on the inter-chromosomal interactions formed by *Prdm1* and uniquely expressed in plasma cells. **(d)** Inter-chromosomal interaction cloud plots showing the positioning of Alzheimer’s genes and ER genes that make inter-chromosomal interactions with *Prdm1* in plasma cells. **(e)** Distribution of the number of *Prdm1* ATAC peaks that interact with the promoters of target genes, including Alzheimer’s genes (left) and ER genes (right) derived from GSEA analysis. *Prdm1* ATAC peaks are defined as ATAC peaks with the *Prdm1* motif (p < 10^−3^).

To validate inter-chromosomal interactions and determine the frequencies of inter-chromosomal interactions in naïve B and differentiated plasma cells, we performed RNA-FISH (**Fig. 7a**). Naïve B and *Blimp*-GFP cells were sorted, formaldehyde-fixed and hybridized with fluorescently labeled intronic *Prdm1*, *Atf4* and *Xbp-1* probes. Consistent with the TCC reads, *Prdm1*, *Atf4* and *Xbp1* inter-chromosomal interactions occurred with high frequency in differentiated plasma cells (**Fig. 7b**). Consistent with transcription, interactions appeared in euchromatic regions depleted of DAPI staining. Finally, we found that inter-chromosomal interactions involving these loci occurred within two hours after activation (**Fig. 7c**). We hypothesized that the purpose of gene co-localization is for synergy of expression. To determine the relationship between Prdm1 and Xbp1 expression, we calculated the Pearson correlation of the intronic spot numbers for these genes. We found a correlation (p-value= 1.21e-06) that supports a synergy of expression between Xbp1 and Prdm1 expression (**Fig. 7d**). Taken together, these data indicate that the differentiation of plasma cells is closely linked with enrichment for a wide spectrum of inter-chromosomal interactions involving members of the UPR pathway (**Fig 7e)**.

**Figure 7.**
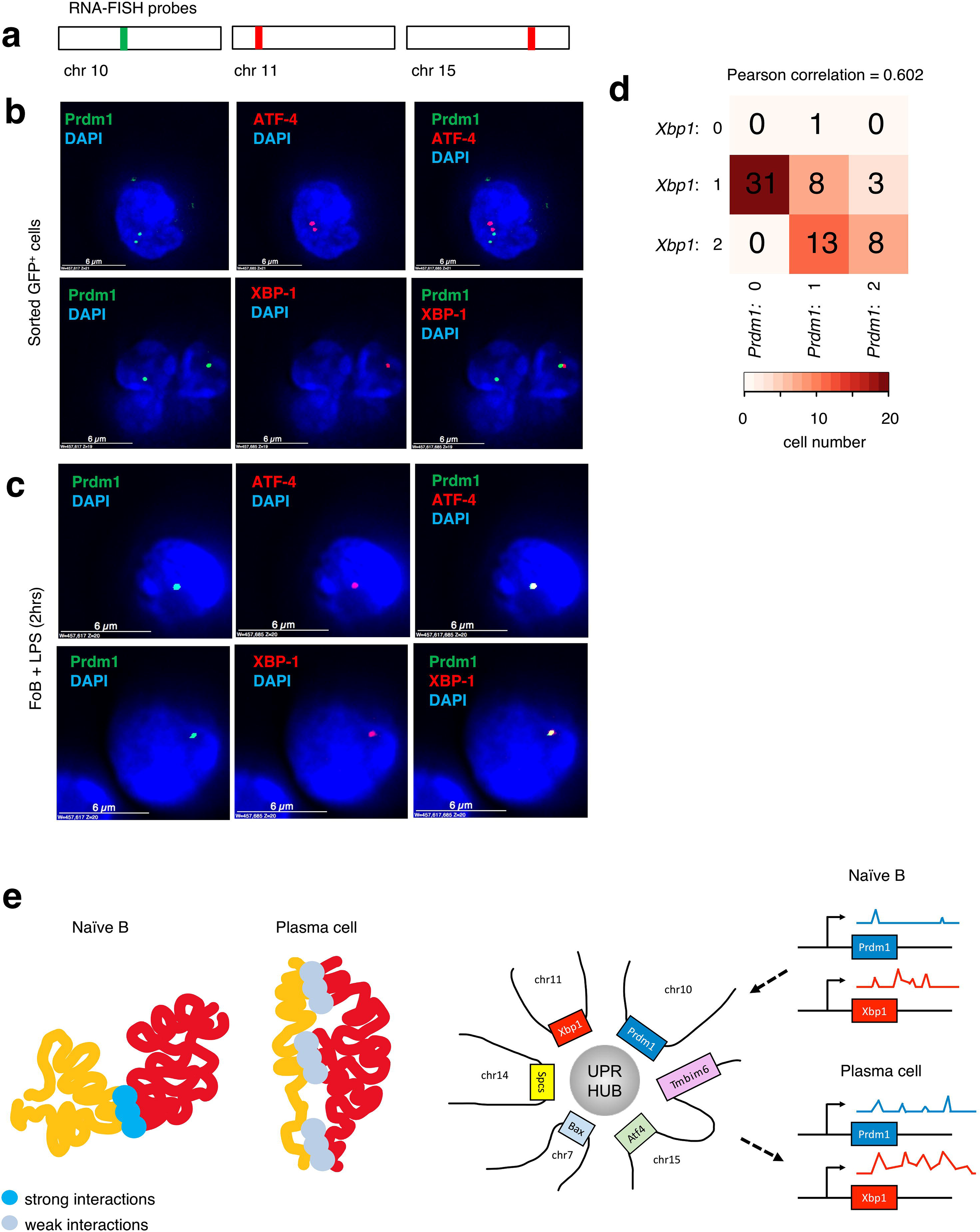
Prdm1 forms inter-chromosomal transcription hubs with ATF4 and XBP1 within hours in LPS-stimulated cells. **(a)** Locations of probe regions used for RNA-FISH experiments. **(b)** Example images of RNA-FISH for *Prdm1* and *Atf4* (top row) or *Prdm1* and *Xbp1* (bottom row) in day 3 sorted GFP^+^ plasma cells**. (c)** RNA-FISH for *Prdm1* and *Atf4* (top row) or *Prdm1* and *Xbp1* (bottom row) in unsorted 2-hour stimulated B cell cultures. **(d)** Heat map displaying the number of cells that express different combinations of intronic RNA-FISH spots. Chi-square test measuring the likelihood that the expression of *Prdm1* and *Xbp1* are independent of each other: p-value=1.21e-06, Pearson correlation=0.601. Colors displaying the most prevalent (darkest) combination to rarest (lightest) combination. **(e)** Model of chromosome confirmation changes during the naïve B to plasma cell transition and its effects on gene expression.

## DISCUSSION

An ensemble of genomic regions switch nuclear location upon establishing B or T cell fate^41^. Prominent examples include cis-regulatory elements associated with the EBF1 and FOXO1 loci that during the transition from the pre-pro-B to the pro-B cell stage reposition from the heterochromatic to the euchromatic compartment^6^. Likewise, upon committing to the T cell lineage, the Bcl11b super-enhancer relocates from the lamina to the euchromatic compartment^7^. Here, we have examined how the genomes of terminally differentiated B cells are assembled. In contrast to early B cell development, the genomes of differentiating plasma cells undergo large-scale decontraction across vast genomic distances. One possible explanation for the observed reduction in long-range intra-chromosomal and decrease in strong inter-chromosomal interactions in plasma cells is the increase in cell size during naïve B cell differentiation. Similar differences were observed between fibroblasts and spermazoa, an example of extreme compactness, where DNA is packed into ~5% the volume of somatic cells^42^.

Plasma cells appeared to lock in a program of gene silencing by repositioning regions inactive in naïve B cells to the heterochromatic compartment to further dampen transcription. Moreover, whereas in developing B cells, the Ebf1 repositioned away from the nuclear lamina, in differentiating plasma cells, the Ebf1 locus switched to a repressive perinucleolar region reminiscent of other silenced early B cell and non-B cell genes during B cell activation^43^. Strikingly, we found that the final stages of plasma cell differentiation are also closely linked with a series of unique structural features, including the adoption of elongated helical chromosomes that collectively fold into a cartwheel-like configuration. We note that α-helical-like chromosomal structures have been observed in sperm cells^44^. Likewise, as early as 1978, Sedat and Manuelidis proposed that interphase chromosomes fold as helical structures^45^. More recently, it was demonstrated that during prometaphase, inner loops are nested within 400kb outer loops that adopt helical structures, which increase in size to span vast genomic distances (>12 Mb)^46^. Radial positioning of chromosomes, similar to the arrangement that we observed in plasma cells, has also been observed in mitotic cells^47^. However, given that plasma cells are non-cycling^48^, we speculate that plasma cells retain the wheel-like rosette of mitotic cells as they fall out of cell cycle into G0/G1^4932^. These observations raise the structure-function question of why plasma cell chromosomes are decontracted and adopt elongated configurations. We suggest that plasma cells adopt a polarized nuclear architecture to facilitate active transport of antibody transcripts, as well as other transcripts closely associated with a plasma cell specific transcription signature, for efficient export to and translation in the expanded endoplasmic reticulum (ER)^19^.

Perhaps the most striking feature of plasma cell nuclear architecture involves the assembly of *de novo* inter-chromosomal interactions. Interphase chromosomes are not randomly distributed but rather occupy distinct regions of the nucleus. In naïve B cells and plasma cells, small chromosomes were positioned predominantly in internal locations whereas large chromosomes localize primarily to peripheral zones, consistent with the size-dependent positioning of chromosomes observed in human interphase nuclei^39^. Although chromosomes fold into distinct structures and rarely intermingle, they are not entirely segregated. Well known examples of inter-chromosomal interactions include the *Ifng, Il4, and Il5* genes in naïve and T helper subsets^50^ and olfactory receptors in sensory neurons^51^.

Here, we identify a novel inter-chromosomal transcription hub that includes *Prdm1, Xbp1* and *Atf4* (**Fig. 7e**). We also observed inter-chromosomal interactions between *Prdm1* and an Alzheimer’s cluster. It is tempting to speculate that the regulatory circuits important for plasma cell metabolism, protein folding, and housekeeping functions may also be dysregulated in Alzheimer’s disease. We also found that, upon B cell activation, the UPR transcription hub assembles in less than two hours. We suggest that the activation of UPR or ER transcription signature requires the coordinate induction of an ensemble of genes, assembled into an inter-chromosomal transcription hub, to collectively activate a plasma cell specific program of gene expression. These findings raise the question of why activated follicular B cells assemble an UPR inter-chromosomal transcription hub with such rapid speed. We suggest that the induction of a plasma cell specific transcription signature requires the simultaneous activation of an enhancer repertoire that is associated with a large spectrum of genes. The assembly of *de novo* inter-chromosomal transcription hubs explains how in differentiating plasma cells groups of genes are simultaneously induced to orchestrate a plasma cell specific program of gene expression.

## AUTHORS CONTRIBUTIONS

A.B. and C.M. conceived of the study and designed experiments. A.B. performed the majority of the experiments and analyzed data. Z.H. analyzed data and designed the computational modeling. M.A. performed 3D-FISH. V.C. designed the modified LMP vector, helped prepare samples, and provided important technical suggestions. M.D. provided analysis tools and important technical suggestions. K.C. and Y.C.L. contributed to analysis. A.B. and C.M. wrote the manuscript. C.M. supervised the study.

## ACKNOWLEDGMENTS

We thank Maho Niwa for critical reading of the manuscript. Sequencing was performed at the Institute for Genomic Medicine Center, University of California, San Diego (F32 GM106631). Imaging was performed at the University of California, San Diego School of Medicine Microscopy Core (NS047101). We thank S. Nutt (Walter and Eliza Hall Institute of Medical Research) for Blimp1-GFP mice, Anjana Rao for the LMP vector, and current and past members of the Murre lab for their advice and assistance throughout the project. A.B. was supported by a Ruth L. Kirschstein National Research Service Award (1F32GM106631-01A1). M.D. was supported by the Frontiers in Innovation Scholars Program. This study was supported by funding from the NIH 4DN Nucleome Project (DK107977), Center for Computational Biology and Bioinformatics (CCBB) (UL1TRR001442) and the NIH to C.M. (AI102853 and DK107977).

## MATERIALS AND METHODS

### Mice

Adult C57BL/6 were purchased from Jackson Laboratories and maintained in a specific pathogen-free facility at the University of California, San Diego, in accordance with institution guidelines for animal care and welfare. B6.Blimp^+/GFP^ were bred and housed in our colony.

### Flow cytometry

Single-cell suspensions of splenocytes were prepared, depleted of red blood cells by hypotonic lysis, and stained with optimal dilutions of the indicated antibodies. All of the following reagents were obtained from eBioscience: anti-CD4 (RM4-5), anti-CD8a (53-6.7), anti-Gr-1 (RB6-8C5), anti-F4/80 (BM8), and anti-TER119; anti-IgD (11-26); anti-B220 (RA3-6B2); anti-CD19 (1D3). Doublets were excluded using the combined width and height parameters of the forward and side scatter parameters. Flow cytometric acquisition was performed on a BD LSRII, and analyses were performed using FlowJo 10.1r5 (Tree Star). Cells were sorted with a three-laser FACsAria Fusion.

### Cell culture

CD23+ follicular B cells were isolated by positive selection using anti-biotin microbeads (Miltenyi Biotec). Sorted B cells were cultured for 3 days in RPMI 1640 medium supplemented with 10% FCS, antibiotics, 2mM _L_-glutamine, and β–mercaptoethanol (50 μM) at 37C, 5% CO_2_, before analysis by flow cytometry. LPS (*Escherichia coli*) was purchased from Sigma-Aldrich; L2654-1MG and used at 10 μg/mL. For cell division experiments, cells were labeled with CellTrace Violet reagent (Molecular Probes). Dead cells were excluded using DAPI (Molecular Probes).

### DNA-FISH

Cells were placed on poly-L-lysine coated coverslips for 30 minutes in 37ºC incubator. fixed with paraformaldehyde at the final concentration of 4% for 10 minutes. The 200 kb bacterial artificial chromosome (BAC) probe RP23-118P17 and 40 kb fosmids was obtained from the BACPAC Resource Center (BPRC) at Children’s Hospital Oakland Research Institute. Probes were labeled by nick translation using Alexa-488 dUTP, Alexa-568 dUTP, or Alexa-647 dUTP (Invitrogen). Incubated overnight at 37ºC. The following morning, coverslips were washed twice in 50% formamide/2x SSC at 37ºC for 30 minutes on a shaking incubator at 300 rpm. Coverslips were then rinsed in 1x PBS containing DAPI. Coverslips were rinsed once more with 1x PBS.

### Immunofluorescence

Coverslips were permeabilized with 0.1% saponin/0.1% Triton-X in PBS at room temperature for 10 minutes, then incubated for 20 minutes at room temperature with 20% glycerol in 1x PBS. Slides were submerged in liquid nitrogen three times, rinsed once with 1x PBS, and then blocked for 30 minutes with 5% BSA/0.1% Triton-X in 1x PBS for 30 minutes at 37ºC. The nucleolus was stained first with primary antibodies to B23 (Abcam) for 30 minutes in a humidified chamber at 37ºC. Coverslips were washed twice, 10 minutes each, in 0.1% Triton-X in 1x PBS at room temperate on a shaker at 70 rpm. Secondary staining with performed using donkey antibody to mouse IgG conjugated to Alexa-568 (Invitrogen) for 30 minutes at 37ºC in 5% BSA/0.1% Triton-X and 5% serum in 1x PBS. Coverslips were washed twice for 10 minutes each in 1x PBS/0.1% Triton-X at room temperature with gentle agitation (70 rpm). The nuclear lamina was stained first with primary antibody to Lamin B1 (sc-6217; Santa Cruz Biotechnology), followed by secondary staining.

### Single molecule RNA-FISH

Pools of fluorescently labeled oligonucleotide probes per RNA were designed and purchased from LGC Biosearch Technologies using the Stellaris Probe Designer. For intronic *Prdm1* and *Xbp1*, default settings were used. For intronic *Atf4*, masking level was set to 3, oligo length was set to 18, and nucleotide spacing was set to 1. Cells were fixed and permeabilized as described for DNA-FISH, but were stored overnight at 4C in 70% ethanol rather than PBS. The next day, coverslips were rinsed in wash buffer (10% formamide in 2x SSC) to remove ethanol, then incubated in wash buffer for 5 minutes at room temperature. Coverslips were incubated with probe of interest in hybridization buffer overnight in a 37C humidified hybridization oven. The next morning, coverslips were rinsed in washed buffer and washed twice for 30 minutes each in 37ºC temperature-controller shaker at 100 rpm. During the second wash, DAPI was added to the wash buffer. Slides were rinsed once in 2x SSC, once in 1x PBS, and then mounted with Prolong Gold Anti-Fade reagent. Slides were dried in the dark for at least 6 hours prior to sealing with nail polish and imaging.

**TABLE:** RNA FISH probes

| Region of interest | Region | Chromosome | Position (bp) | # probes | Fluorophore |
| --- | --- | --- | --- | --- | --- |
| Intronic Prdm1 | Fourth intron | Chromosome 10 | 44161971-44166597 | 48 | Quasar 670 |
| Intronic Xbp1 | First two introns | Chromosome 11 | 5421204-5421925;<br>5422023-5423335 | 36 | Cal Fluor 610 |
| Intronic Atf4 | All introns (2) | Chromosome 15 | 80086116-80086636;<br>80086952-80087063 | 22 | Cal Fluor 610 |

### Imaging

Three-dimensional fluorescent images were acquired using an Applied Precision Inverted Deconvolution Deltavision Microscope with a 100x objective (Nikon 100x/1.40 oil or Olympus 100x/1.4 oil). Optical sections (z-stacks) 0.2µm apart were obtained throughout the cell volume in the DAPI, FITC, Red, and Cy5 channels. Deconvolution was performed using Softworx Software version 5.0 with the following settings: 10 cycles of enhanced ratio (aggressive), camera intensity offset = 50, normalize intensity and apply correction options selected.

### Calculating distances between DNA loci

The nuclei of individual cells were identified by DAPI staining, and cells containing two spots per DNA-FISH or intronic RNA-FISH channel were verified manually. Images were cropped to contain single cells. Analysis of cells in three dimensions was performed using the FIJI plugin Tools for Analysis of Nuclear Genome Organisation (TANGO) version 0.94^52^. Distance between two DNA-FISH spots was calculated after segmentation in TANGO by running the “Distance” measurement between center-of-mass of the two loci. Distance between nuclear structures and a DNA-FISH spot was calculated as the minimum distance between the center-of-mass of the DNA-FISH spot to the edge of the corresponding structure. Negative values indicate inclusion of the FISH spot in the nuclear structure.

### RNA-Seq sample preparation

A total of 10,000-100,000 B cell subsets were sorted into Buffer RLT (QIAGEN) and RNA was isolated. Total RNA was isolated using RNeasy Mini kit (Qiagen) with on column DNase treatment. RNA was again treated with TURBO DNase (Life Technologies) and mRNA was purified with Dynabeads mRNA purification kit (Life Technologies. First-strand synthesis kit (Life Technologies) in presence of actinomycin D using a combination of random hexamers and oligo(dT). Second-strand synthesis was performed with dUTP instead of dTTP. The ds-cDNA was sonicated to 150-400 bp using the Covaris sonicator. Sonicated cDNA was ligated to adaptors. The resulting DNA was treated with uracil-N-glycosylase prior to PCR amplification with the indexing primers. Following PCR, fragments were size-selected and sequenced. Single-end sequencing reads were mapped to mouse genome build mm9 using TopHat and analyzed using the Cufflink-cuffdiff pipeline.

### Analysis of RNA-Seq

Single-end sequencing reads were mapped to mouse genome build mm9 using TopHat and converted to track plot by IG^53,54^. Transcript quantification was performed using Kallisto with default settings. Modified TPM was used in cross-sample comparison, as plasma cells are known to have more RNA than naïve B cells. Plasma TPM was multiplied by 2.1 to account for this difference. Fold change was computed by calculating fold changes between TPM+1.

### Overexpression

The GFP cassette of the LMP vector was replaced with hCD25 cassette for EBF1 overexpression. Retroviral supernatant was obtained through 293T transfection by the calcium phosphate method with these constructs in conjunction with the packaging plasmid pCL-Eco. CD23^+^ cells were MACs selected and stimulated in vitro for 48 hours prior to infection. Cells were harvested 48 hours after infection for analysis by flow cytometry.

### TCC

TCC was performed as previously described^35^ using 5 million naïve B cells or sorted plasma cells.

### Analysis of TCC data

TCC reads were mapped by bowtie and processed by HOMER to create tag directory with default setting^55^. Read number of naïve and plasma cells were down-sampled by HOMER, using getRandomReads.pl and reducing the total number of reads to match the smaller sample size (naïve B cells). Inter-chromosomal interactions were called by HOMER using 1 Mb bins and a threshold p=10^−4^ with the other options set as default. Intra-chromosomal interactions were called by HOMER using 25 kb bins, threshold p=10^−4^ and max distance 3 Mb with the other options set as default. PC1 and corrDiff were calculated by HOMER with 25 kb, 50 kb, 1 Mb, and 2 Mb resolution. TCC matrices, normalized or raw, were generated by HOMER with default setting using 1Mb bins for the whole genome and 50 kb bins for individual chromosomes. The raw matrices were normalized by KR-normalization^56^.

#### Compartments

To quantify switched domains, the following criteria were used: regions with PC1 change more than 25, regardless of sign, were considered switched. Bin size was 50 kb. Consecutive regions were merged. Two regions separated by only one bin that did not pass threshold were also merged.

#### corrDiff

As explained by HOMER: “PCA analysis is a useful tool for analyzing single TCC experiments. However, comparing PC1 values between two experiments is a little dangerous. PCA is an unbiased analysis of data, and while PC1 values usually correlated with "active" vs. "inactive" compartments, the precise qualitative nature of this association may differ slightly between experiments. Because of this, it is recommended to directly compare the interaction profiles between experiments rather than simply looking at PC1 values.

One way to do this is to directly correlate the interaction profile of a locus in one experiment to the interaction profile of that same locus in another experiment. If the locus tends to interact with similar regions in both TCC experiments, the correlation will be high. If the locus interacts with different regions in the two experiments, the correlation will be low. Note that this is NOT PCA analysis, but a direct comparison between two experiments at each locus. By default, this will compare the interaction profiles across each chromosome (inter-chromosomal interactions are generally too sparse for this type of analysis).”

#### Average intra-chromosomal contact maps

The average intra-chromosomal TCC contact maps for naïve B cells and plasma cells (**Fig. 1**) were generated as follows: 1) calculate the 100Kb resolution raw TCC contact maps for each chromosome (chr1-19 and X, in total 20 matrices); 2) resize each contact matrix to have the 100 rows and 100 columns; and, 3) calculate the average contact intensity of all chromosomes.

#### Average inter-chromosomal contact maps

The average inter-chromosomal TCC contact maps for naïve B cells and plasma cells (**Fig. 5**) were generated as follows: 1) calculate the 100 kb resolution raw TCC contact maps for each pair of chromosome (chr1-chr2, chr1-chr3 … chrX-chr19, in total 190 matrices); 2) resize each contact matrix to have the 100 rows and 100 columns; 3) calculate the average contact intensity of all chromosome pairs.

### Modeling

In order to model the average chromosomal configuration from TCC data, we need to map each TCC bin from a high dimensional space to a low dimensional space (i.e., 3D or 2D). The dimension of TCC matrices is determined by the number of rows (bins). These models provide a more intuitive view of the TCC data. The modeling of **Fig. 4** was performed as follows:

1) Convert the KR-normalized whole genome matrix to a correlation matrix and then calculate a distance matrix using 1 minus the correlation matrix. There are at least two inherent limitations when modeling chromosome configuration using TCC data: (A) the read number is not an accurate measurement of the average spatial distance between two loci, but rather a measurement of the probability that two loci co-localize; and, (B) the read number is effected by various batch effects (i.e., technical preparation of the library) and subject to sampling noise (i.e., sequencing bias). Converting the TCC matrix to a 1 minus correlation distance matrix partially offsets these problems by narrowing the possible range of distances between two bins to 0 to 2 rather than 0 to the max number of reads per bin. For example, if two loci are in spatial proximity (i.e., in the same chromosomal territory) of one another but do not interact, then 0 reads between the two loci would be detected. Thus, the predicted spatial distance based on these TCC reads would be very far apart. This predicted distance does not match the actual situation. Therefore, a distance conversion must be performed to reduce the “penalty” of low read numbers between bins (corrects for limitation A above). The conversion also corrects for some experimental noise as the correlation of interaction patterns are more stable than read numbers (corrects for limitation B above).

2) Apply t-SNE to map the loci to 3D or 2D space, according to the distance matrix generated in step 1. t-SNE is a non-linear dimensionality reduction algorithm^57^. Consider each region as a dot, and their distance to each other is indicated by the distance matrix generated in step 1. t-SNE will try to find the best position of these dots in a given space subject to the limitation of the distance matrix. Additionally, t-SNE focuses on preserving short-range rather than the long-range distances. This feature also helps to resolve limitation A discussed in step 1. The Rtsne package was employed^58^. With 1Mb bins, t-SNE perplexity was set to 75 (perplexity 60 to 125 can produce similar results). Because of the inherent randomness of t-SNE, each run of the simulation will produce slightly different results but the overall conformations for naïve B cells and plasma cells remain the same.

For the inter-chromosomal cloud plots in **Fig. 6**, only the inter-chromosomal significant interactions with p<10^−4^ were considered. The 1Mb region bins linked by significant interactions were assigned an arbitrary distance of −1; the regions not connected had an arbitrary distance of 0. t-SNE was employed to map the interaction relationship between these bins to 2D. Perplexity was set to 30 (perplexity in 20-50 result in similar results).

### ATAC-seq

ATAC-seq was performed as previously described^59^. 50,000 cells were used for library preparation. Cells were treated with transposition mix for 30 minutes at 37ºC. DNA was purified with DNA Clean & Concentrator (Zymo Research). Library fragments were amplified by PCR using 1x NEBnext PCR master mix and 1.25 uM of custom Nextera PCR primers 1 and 2 as follows: 72ºC for 5 min., 98ºC for 30 seconds, 98ºC for 10 seconds, 63ºC for 30 seconds and 72ºC for 1 minute. Amplified libraries were selected using SPRI beads. The quality of the library was measured using Agilent TapeStation and the final product was sequenced on an Illumina Hi-Seq4000.

### Analysis of ATAC-seq and ChIP-seq

Reads were mapped to mouse genome build mm9 using TopHat and converted to track plots by IGV. ATAC peaks were called by MACS 1.4 with default settings for broad peak. ATAC peaks were associated with promoters of genes based on the TCC interactions. If one anchor of a significant interaction overlapped with the promoter of the target gene and the other anchor overlapped an ATAC peak, then the ATAC peak was considered to interact with the promoter with a p-value equal to the TCC interaction. PRDM1 motifs within ATAC were called by FIMO^60^ with markov 8 background and the mouse PRDM1 motif derived from cisBP^61^.

### GSEA Analysis

Significant inter-chromosomal interactions were identified using HOMER at the p-values noted in the text. Interactions originating from *Prdm1* were selected and annotated with all the genes overlapping the 1 Mb bins. Genes unique to plasma cells were interrogated in GSEA by computing overlaps with available gene sets. The enrichment of these annotations was calculated as a ratio: the fraction of genes that match a given term to the fraction of all measured genes with the given term. The p-value for this enrichment was calculated using the hypergeometric function.

